# PlasChain: an algorithm for improving long plasmid reconstruction from metagenome assemblies

**DOI:** 10.64898/2026.07.27.740866

**Authors:** Shi Feng, Haonan Song, Ron Shamir, Lianrong Pu

## Abstract

**Background:** Plasmids play a critical role in horizontal gene transfer and the spread of antibiotic resistance. However, recovering complete plasmid sequences from metagenomic samples remains highly challenging due to extensive repeat content, structural heterogeneity, and large variation in plasmid size. Existing methods typically identify plasmids from metagenome assemblies by exploiting coverage differences or detecting minimum-weight cycles in assembly graphs. While effective for dominant plasmids, these approaches often fail to recover low-abundance and long plasmids.

**Results:** Here we present PlasChain, a novel algorithm designed to improve plasmid assembly and identification from complex metagenomic data. Building upon the cycle-peeling strategy of SCAPP, PlasChain incorporates contig path information and a cycle-merging procedure to prevent long plasmids from being fragmented into multiple shorter cycles. In addition, PlasChain jointly leverages paired-end read alignments, sequence composition patterns, and coverage variation to filter out assembly artifacts and reduce false positives. We evaluated PlasChain against state-of-the-art plasmid assemblers, including SCAPP and metaplasmidSPAdes, using a diverse set of simulated and real metagenomic datasets. Across nearly all benchmarks, PlasChain demonstrates superior performance in recovering long plasmids while maintaining competitive accuracy in assembling short plasmids. Furthermore, analysis of real metagenomic samples shows that PlasChain is capable of assembling previously uncharacterized plasmids, including putative megaplasmids that are typically underrepresented in current plasmid databases.

**Conclusions:** PlasChain is a novel graph-based plasmid assembler that improves the recovery of long plasmids from short-read metagenomic data. Evaluation on diverse simulated and real metagenomic datasets demonstrates that PlasChain consistently outperforms existing plasmid assemblers, especially for long plasmid reconstruction. These results highlight the potential of PlasChain to facilitate comprehensive characterization of plasmid diversity, antimicrobial resistance, and horizontal gene transfer in complex microbial communities. The source code and testing data of PlasChain are freely available at https://github.com/SDU-ACG-Lab/PlasChain.

**Supplementary Information:** This manuscript is accompanied by a supplementary file.

## Background

Plasmids are self-replicating genetic elements that can transfer between bacterial hosts, playing a crucial role in horizontal gene transfer (HGT). They facilitate the spread of traits such as antimicrobial resistance and virulence [10, 6], making them central to the emergence of multidrug-resistant pathogens and posing a growing threat to public health. As a result, there has been increasing interest in plasmid detection and characterization, not only to understand their roles in antibiotic resistance but also to explore their potential applications in biotechnology and clinical settings.

Traditionally, biologists isolate a single bacterial strain and cultivate it in the laboratory to obtain and sequence a DNA-rich sample specific to that strain. Tools such as plasmidSPAdes [1] and Unicycler [26] were developed to assemble plasmids from these isolates. Since such isolated samples contain only a single bacterial strain, differences in contig coverage can be leveraged to efficiently distinguish plasmids from chromosomes, as plasmids typically exhibit a higher copy number. For instance, plasmidSPAdes iteratively filters out chromosomal contigs and simplifies the assembly graph based on coverage differences, while Unicycler integrates the accuracy of short reads with the structural resolution of long reads, utilizing both contig coverage and the topological structure of the assembly graph to resolve plasmid paths.

Although these tools are effective for plasmid assembly from isolated strains, many organisms remain unculturable in laboratory settings [12, 21]. Advances in high-throughput sequencing have led to the widespread availability of large-scale metagenomic datasets [3], providing new opportunities to explore plasmid diversity directly from environmental samples.

However, assembling complete plasmids from metagenomic data presents significant challenges due to the complexity of species composition and wide variation in abundance. This variability limits the effectiveness of tools originally designed for isolated strains [16]. For example, high-coverage contigs in metagenome assemblies may represent plasmids from rare species with high copy numbers or chromosomes from more abundant species.

To address these challenges, metaplasmidSPAdes [2] (abbreviated henceforth to mpSPAdes) and SCAPP [18] were developed to assemble complete plasmids from metagenomic sequencing data. mpSPAdes builds upon metaSPAdes [4] by iteratively removing low-coverage edges, progressively increasing the coverage cutoff at each iteration in order to simplify the assembly graph [2]. Plasmids within these subgraphs are detected using ExSPAnder [20], and further verified by plasmidVerify [2]. This iterative refinement allows mpSPAdes to improve upon plasmidSPAdes by resolving dominant plasmids better. However, it still struggles to recover low-coverage plasmids that are entangled with chromosomal edges. SCAPP builds upon the concept of peeling cycles from the assembly graphs introduced by Recycler [22]. It incorporates biological features of plasmids, such as plasmid-specific genes (PSGs), and a plasmid likelihood score generated by plasmid classifiers, to assign weights to edges in the assembly graph. This way, edges likely originating from plasmids receive lower weights. SCAPP then assembles complete plasmid sequences by identifying cycles with small edge weight from the assembly graph. While SCAPP achieves high precision in assembling short plasmids, its greedy strategy of detecting minimum-weight cycles limits its effectiveness when dealing with long or linear plasmids.

To overcome these limitations, here we present PlasChain, an algorithm specifically designed to enhance the recovery of long plasmids (≥ 10kb) from metagenomic assemblies. Building upon the SCAPP framework, PlasChain incorporates contig path information from metaSPAdes alongside supporting evidence to identify long plasmids. In situations where contig path information is unavailable, PlasChain employs an additional cycle-merging step to detect and integrate cycles that are likely derived from the same plasmid. Evaluation on both simulated and real datasets demonstrates that PlasChain outperforms SCAPP and mpSPAdes in most scenarios, particularly in the recovery of long plasmids.

## Methods

### Overview of the PlasChain algorithm

PlasChain accepts as input a set *R* of paired-end reads and an assembly graph *G* = (*V, E*) assembled from *R* by metaSPAdes. In the assembly graph, each node in *V* represents a linear sequence, while each directed edge (*u, v*) in *E* represents an overlap between the suffix of *u* and the prefix of *v. u* and *v* are also called the *endpoints* of the that edge. Building on the SCAPP framework, PlasChain identifies plasmids as cycles exhibiting uniform coverage within the assembly graph. The algorithm begins by identifying high-confidence plasmidic nodes based on the presence of PSGs or confident prediction by the plasmid classifier PlasClass. Then, minimum-weight cycles that pass through these high-confidence plasmidic nodes and exhibit uniform coverage are iteratively identified and peeled from the graph. The cycle-peeling process continues until no cycles meet the predefined criteria. Details on the weight assignment of nodes and the cycle-peeling process are provided in the following subsections. A key innovation of PlasChain lies in its ability to identify long plasmids, particularly when supported by contig path information. In cases where such support is lacking, PlasChain applies an additional cycle-merging step to detect and combine cycles that are likely to originate from the same plasmid. An overview of the PlasChain algorithm is provided in Figure 1 and 2.

**Fig. 1.**
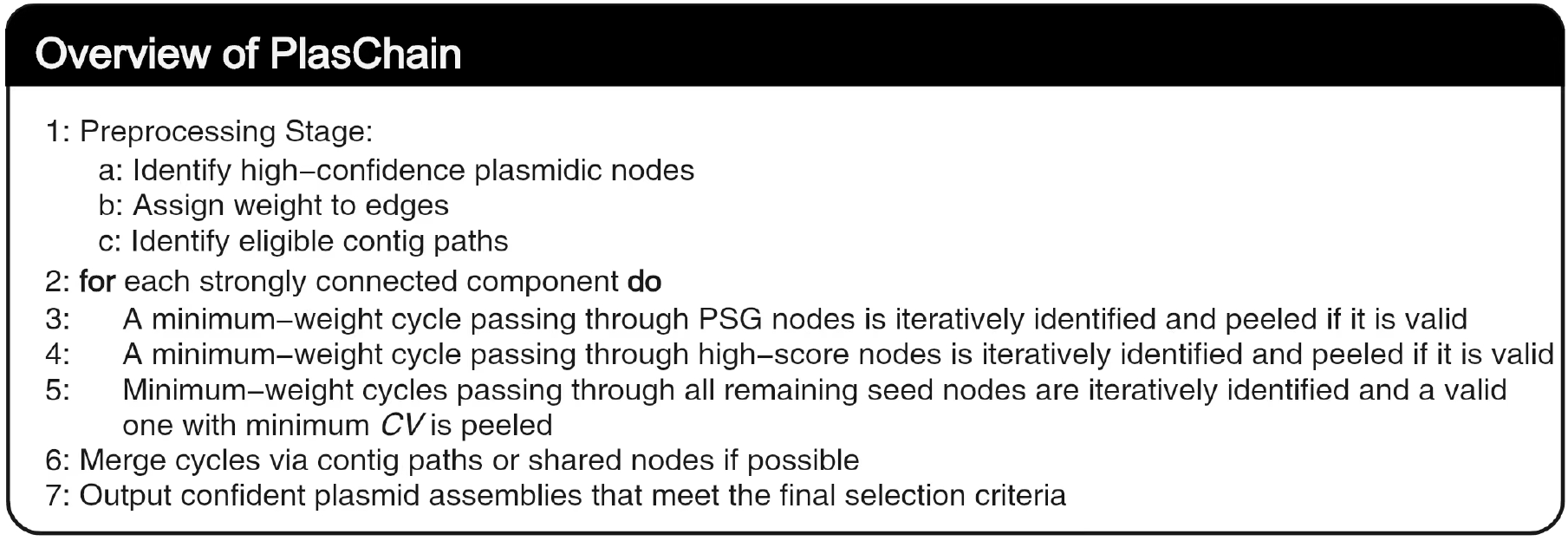
Overview of the PlasChain Algorithm.

**Fig. 2.**
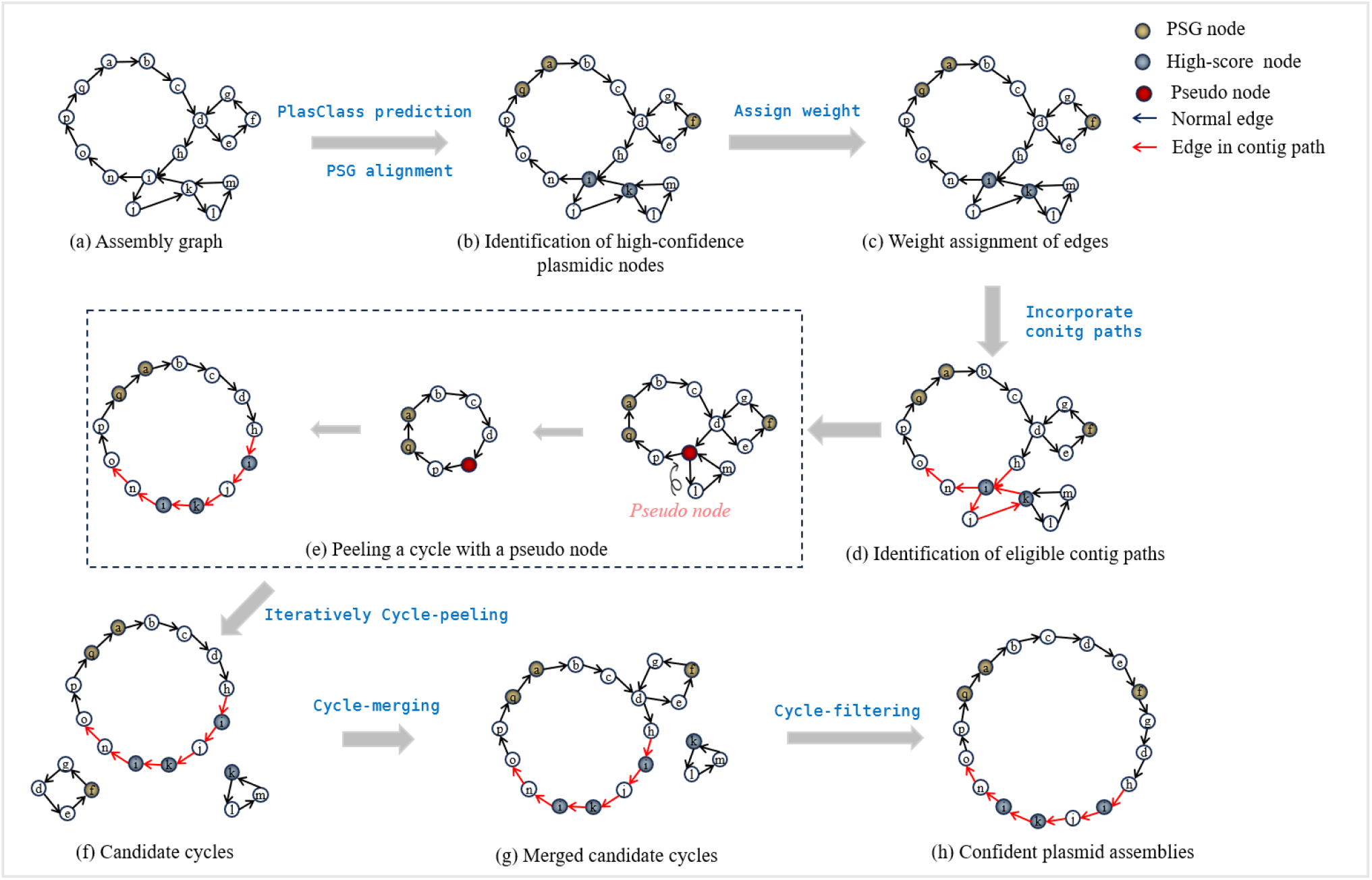
A graphic overview of PlasChain.

### Identification of high-confidence plasmidic nodes

PlasChain begins by identifying nodes in the assembly graph with a high likelihood of plasmid origin. First, all nodes are mapped against a predefined PSG set provided by SCAPP. Nodes with a BLAST match to any PSG sequence exhibiting at least 75% sequence identity and 75% alignment coverage are annotated as *PSG nodes*. Additionally, PlasChain assigns a plasmid probability score *p* (where 0 ≤ *p* ≤ 1) to each node using PlasClass [17], a machine learning-based classifier that estimates the likelihood of plasmid origin of a sequence. While PlasClass performs well for longer sequences, its predictions for shorter sequences are less reliable. To account for this, PlasChain adjusts the plasmid score based on the node length *L* using the formula:

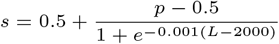

Nodes with an adjusted plasmid score *s >* 0.9 and length greater than 10kb are classified as *high-score nodes*. High-confidence plasmidic nodes are defined as those that are either high-score nodes or PSG nodes. These nodes are prioritized during the subsequent cycle-peeling step. Furthermore, to simplify the assembly graph, nodes with an adjusted plasmid score *s <* 0.2 and length *>* 10kb are removed, as they are assumed to be chromosomal sequences.

### Weight assignment to edges

Before applying the cycle-peeling strategy for plasmid assembly, PlasChain assigns weights to edges in the assembly graph. Edges with lower weights are considered more likely to originate from plasmids and are prioritized for peeling in the subsequent step. In contrast to SCAPP, which assigns edge weights based on the scores of one endpoint, PlasChain incorporates additional factors in its edge weight calculation, including tetranucleotide composition similarity, discounted coverage similarity, and mate-pair read support between the two endpoints. This weight assignment approach allows PlasChain to assess whether adjacent nodes are likely to originate from the same plasmid. Specifically, for an edge (*u, v*), its weight *w*(*u, v*) is defined as:

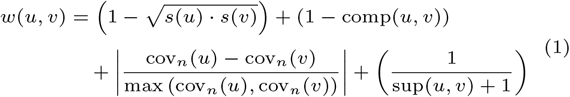

Here, *s*(*u*) and *s*(*v*) denote the length-adjusted plasmid scores of nodes *u* and *v*, respectively. *comp*(*u, v*) represents the cosine similarity of the tetranucleotide composition between nodes *u* and *v*, while *sup*(*u, v*) is the number of mate-pair reads that support the edge (*u, v*). An edge (*u, v*) is supported by a mate-pair read if one of its mates is mapped to node *u* and the other is mapped to node *v*. The terms *cov*_*n*_(*u*) and *cov*_*n*_(*v*) represent the normalized coverages of nodes *u* and *v* with respect to edge (*u, v*), and are computed as:

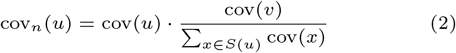

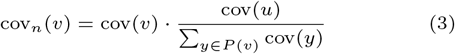

where *cov*(*u*) and *cov*(*v*) represent the original coverage of nodes *u* and *v*, respectively, as generated by the assembler. *S*(*u*) and *P* (*v*) denote the set of successors of *u* and the set of predecessors of *v*, respectively. The use of normalized coverage enables PlasChain to better capture the coverage similarity between endpoints, as nodes may participate in multiple paths or cycles. In summary, PlasChain prioritizes edges that exhibit high plasmid scores, strong similarity between their endpoints, and substantial mate-pair read support, leading to more accurate plasmid assembly.

### Identification of eligible contig paths

Recall that each contig generated by metaSPAdes is represented as either a single node or a path within the assembly graph. These contig paths typically correspond to subpaths of an underlying genome and therefore offer valuable long-range connectivity information for assembling long plasmids [13] (see Figure 3). In this step, PlasChain improves long plasmid recovery by performing three procedures: trimming dead-end nodes from contig paths, identifying plasmid-like contig paths, and compressing eligible contig paths into pseudo-nodes.

**Fig. 3.**
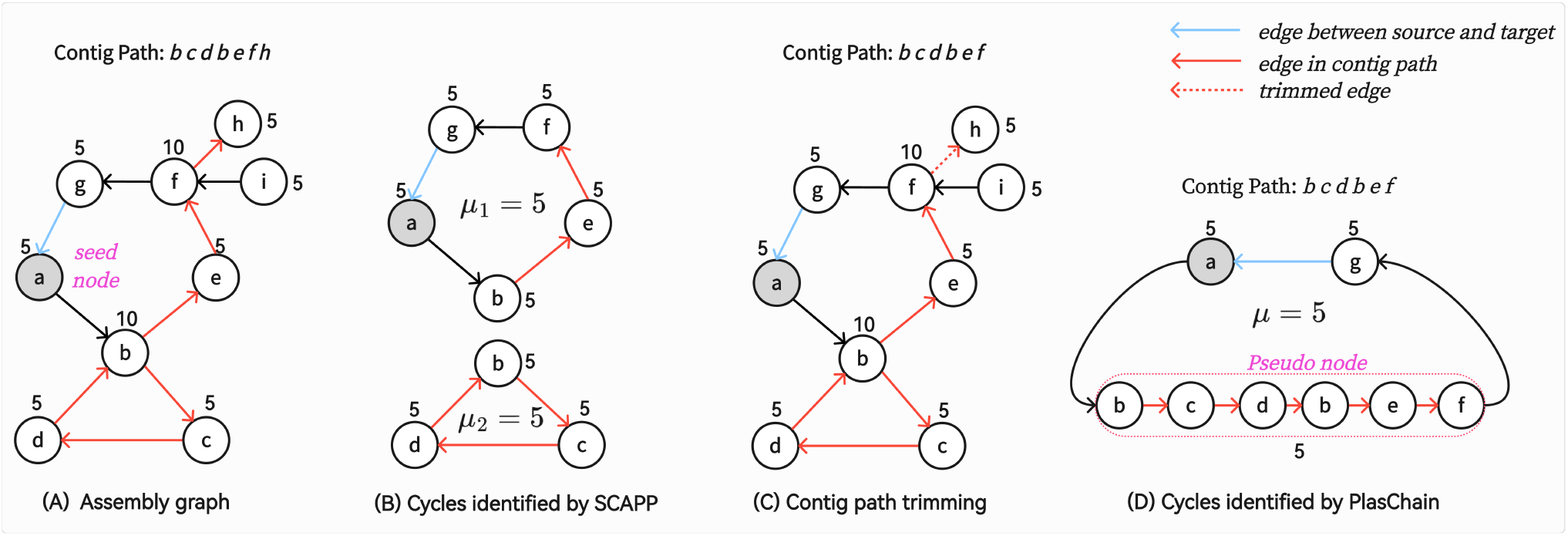
The detection of a long plasmid by using contig path information. Numbers near the nodes indicate their coverage, and all nodes have the same length. (A) An assembly graph with a contig path ‘bcdbefh’. When searching for plasmid-like cycles passing through seed node *a*, Dijkstra’s algorithm takes nodes *a* and *g* as source and target nodes, respectively, and aims to identify a shortest path from source to target. (B) SCAPP fragments the long plasmid into two shorter cycles by searching for cycles of minimum weight. (C) PlasChain removes dead-end node ‘h’ from the contig path. (D) PlasChain successfully recovers the complete plasmid by compressing the contig path into a pseudo-node.

We first remove dead-end nodes from the paths. A dead-end node is defined as a node with either an in-degree or out-degree of zero. Since contig paths originating from circular plasmids are expected to lie within cycles in the assembly graph, they are unlikely to contain dead-end nodes. Thus, PlasChain iteratively trims dead-end nodes from both ends of each contig path until no such nodes remain (see Figure 3(C)).

After trimming, we apply PlasClass to all processed contig paths, retaining only those with a predicted plasmid probability *p >*0.5 and length exceeding 10 kb for further analysis. Contig paths with *p* ≤0.5 are more likely to originate from chromosomal sequences and are therefore excluded. By default, PlasChain further requires that each retained contig path spans at least four distinct nodes. Contig paths meeting these criteria are considered *eligible* and are subsequently compressed into pseudo-nodes. When assigning weight to an edge (*x, v*) connecting a pseudo-node *x* derived from path *P* = *p*_1_, *p*_2_, …, *p*_*k*_ to another node *v*, the length-adjusted plasmid score and tetranucleotide composition of node *x* are calculated using the concatenated sequence of all nodes in *P* . The coverage *cov*(*x*) is defined as the length-weighted mean coverage across all nodes in *P* . Mate-pair support between *x* and *v* is aggregated from all nodes in the path as:

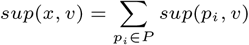

Figure 3 illustrates an example in which SCAPP fragments a long plasmid into two shorter cycles, whereas PlasChain successfully recovers the complete plasmid by incorporating contig path information.

### Priority-based cycle-peeling

PlasChain aims to assemble plasmids by iteratively peeling cycles with uniform coverage from the assembly graph. Specifically, it processes each strongly connected component independently, selecting seed nodes sequentially from three categories: (1) PSG nodes, (2) high-score nodes, and (3) pseudo-nodes derived from eligible contig paths or remaining nodes longer than 1 kb. Within each category, seed nodes are processed in decreasing order of *C* ∗ *L*, where *C* and *L* represent the coverage and length of the seed nodes, respectively. This prioritization ensures that nodes with higher coverage and longer length, which are typically more reliable, are processed first.

For each seed node *v*, PlasChain identifies a minimum-weight cycle passing through *v* using a priority-aware variant of Dijkstra’s algorithm that prioritizes traversal along contig paths. This integration is implemented as follows:

- **Substitution of a seed node with pseudo-node:** If the seed node *v* lies within an eligible contig path *P* = *p*_1_, *p*_2_, …, *v*, …, *p*_*n*_, it is replaced by a pseudo-node *x* representing the entire contig path *P* . Once the minimum weight cycle *C* = *c*_1_, *c*_2_, …, *x* passing through *x* is identified by Dijkstra’s algorithm, PlasChain substitutes *x* back with *P*, yielding a complete cycle *C* = *c*_1_, *c*_2_, …, *p*_1_, *p*_2_, …, *v*, …, *p*_*n*_.
- **Preference for pseudo-nodes during path extension:** During the path extension step of Dijkstra’s algorithm, PlasChain prioritizes pseudo-nodes whenever available. This modification helps prevent fragmentation of long plasmids into several shorter cycles, particularly when the plasmid is supported by high-confidence contig paths.

To evaluate whether a cycle detected above represents a potential plasmid, PlasChain aligns the input read set *R* to the nodes of the assembly graph. A node *v* within a cycle *C* is defined as *mate-pair consistent* if at least half of the reads mapped to *v* have both mates aligned to nodes within *C*. A cycle is considered mate-pair consistent if at least half of its nodes are mate-pair consistent. For cycles consisting of a single node (i.e., self-loops), mate-pair consistency is defined more strictly: ≥ 90% of the reads mapped to the node must have both mates aligned to it.

To assess coverage uniformity, PlasChain computes the coefficient of variation (*CV* ) for each cycle *C* as *CV* (*C*) = *sd*_*cov*′_ (*C*)*/µ*_*cov*′_ (*C*), where *sd*_*cov*′_ (*C*) and *µ*_*cov*′_ (*C*) denote the length-weighted standard deviation and mean of the discounted coverage across all nodes in *C*, respectively. Here, we utilize discounted coverage, as described in SCAPP [18]. The definition of discounted coverage is provided in the Supplementary Materials.

A cycle *C* is considered *valid* if it is a self-loop or it is mate-pair consistent and *CV* (*C*) ≤ 0.5. For seed nodes that are PSG nodes or high-score nodes, once a cycle with minimum weight passing through the node is detected and it is valid, PlasChain immediately peels off the cycle and updates the assembly graph. Cycles are peeled in the order of seed node priority: first PSG nodes, then high-score nodes. For all remaining seed nodes, PlasChain computes the minimum-weight cycle passing through each seed node simultaneously. Among these cycles, PlasChain selects and peels off a valid cycle with the minimum *CV* value, giving preference to cycles that include pseudo-nodes. After peeling the cycle, PlasChain updates the assembly graph and repeats the process of cycle detection and selection of the best valid cycle to peel. This iterative process continues until no valid cycles remain, indicating that all plausible plasmid-like structures have been exhaustively extracted from the graph. All valid cycles peeled from this step are referred to as *candidate cycles*, which are then subjected to further verification in the subsequent steps.

### Cycle merging

Although PlasChain incorporates contig path information to peel long cycles from the assembly graph, the resulting candidate cycles may still represent only partial plasmid genomes rather than complete ones. This is particularly common for plasmids containing repeat sequences, which are often represented in the assembly graph as nodes shared by multiple cycles. Such cycles may be independently peeled from the graph, resulting in fragmented plasmid sequences (see Figure 4). To reconstruct full-length plasmids from these fragmented candidates, PlasChain applies a cycle merging strategy as follows.

**Fig. 4.**
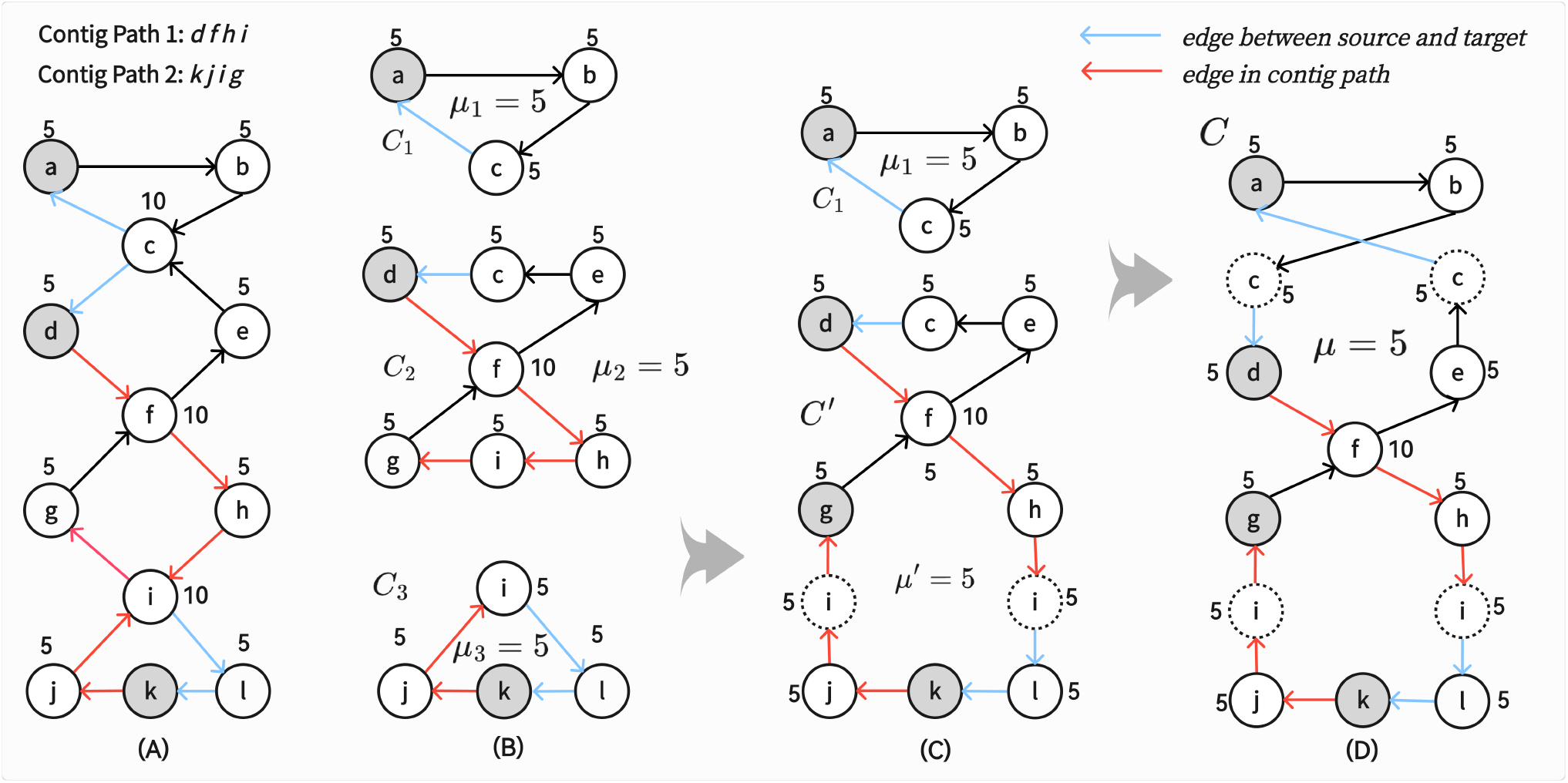
The detection of long plasmids by the cycle-merging step. Numbers near the nodes indicate their coverage, and we assume all of them have the same length. (A) An assembly graph with two contig paths ‘dfhi’ and ‘kjig’; (B) PlasChain peels off three candidate cycles *C*_1_, *C*_2_, and *C*_3_ from the assembly graph; (C) PlasChain merges candidate cycles *C*_2_ and *C*_3_ according to contig path 2; (D) Two cycles in (C) are further merged through the shared node c and forming a long cycle ‘abcdfhilkjigfec’.

Two candidate cycles, *C*_1_ and *C*_2_, are considered for merging only if they share at least one node. Additionally, PlasChain checks whether their coverage and tetranucleotide composition are sufficiently similar according to the following criteria:

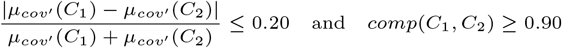

where *µ*_*cov*′_ (*C*_1_) and *µ*_*cov*′_ (*C*_2_) denote the length-weighted mean discounted coverage of cycles *C*_1_ and *C*_2_, respectively, and *comp*(*C*_1_, *C*_2_) represents the cosine similarity between their tetranucleotide composition vectors. The thresholds of 0.2 and 0.9 were determined empirically using validation datasets. Details are provided in the Supplementary Materials.

If these criteria are satisfied, PlasChain attempts to merge the two cycles through either contig path information or shared nodes.

- **Merging via contig path**. All shared nodes between *C*_1_ and *C*_2_ are examined in descending order of coverage. If a shared node *s* lies on a contig path, and there exists a subpath (*a, s, b*) such that *a* ∈ *C*_1_ and *b* ∈ *C*_2_, the two cycles are merged through node *s*. An illustrative example is shown in Figure 4 (C).
- **Merging via a shared node**. If no suitable contig path supports the merge, PlasChain selects the shared node with the highest coverage, denoted *s*_*max*_, and merges *C*_1_ and *C*_2_ through *s*_*max*_. An example is shown in Figure 4 (D).

To avoid false positives, PlasChain merges *C*_1_ and *C*_2_ into a new cycle *C* only if *C* is a valid cycle and its plasmid score exceeds 0.5. If the merge is performed via a shared node rather than a contig path, we further require that the plasmid score of *C* is higher than that of both *C*_1_ and *C*_2_. Only under these constraints is the merged cycle *C* added to the candidate cycle set, and the original cycles *C*_1_ and *C*_2_ are removed. This merging procedure is applied iteratively to the updated set of candidate cycles until no further merges can be performed.

The cycle-merging step raises the risk to merge distinct plasmids that share repetitive sequences. To test this possibility, experiments on five validation datasets showed that it improved the overall F1 score by 0.6-4.6 percentage points (Supplementary Figure S1), indicating that the strategy yields an overall improvement in plasmid recovery.

### Selection of confident plasmid assemblies

After cycle-peeling and cycle-merging, PlasChain selects high-confidence plasmid assemblies from the set of candidate cycles if they satisfy at least one of the following criteria: (1) The cycle contains at least one PSG node and has a plasmid score *>*0.5. (2) The cycle is an isolated self-loop longer than 1kb and is either a PSG node or has a plasmid score *>* 0.9. (3) The cycle is a mate-pair consistent self-loop longer than 1kb and is either a PSG node or has a plasmid score *>* 0.9. (4) The cycle is classified as plasmidic by the plasmid classifier Platon [23] and has a plasmid score *>*0.5. Here, Platon is used as a complementary tool to PlasClass in the final selection step, as it has demonstrated strong performance in plasmid classification.

## Results

To assess the performance of PlasChain, we benchmarked it against two state-of-the-art plasmid assemblers: SCAPP and mpSPAdes. Our evaluation included seven simulated metagenomes with known reference genomes, and six real metagenomic datasets containing unknown genomes.

### Evaluation metrics

We evaluated PlasChain, SCAPP, and mpSPAdes using precision, recall, and F1 score:

- **Precision** = *TP/*(*TP* + *FP* ): The fraction of correctly assembled plasmids among all assembled plasmids.
- **Recall** = *TP/*(*TP* + *FN* ): The fraction of correctly assembled plasmids among all reference plasmids.
- **F1 score** = 2 ∗ *precision* ∗ *recall/*(*precision* + *recall*).

Here, *TP* (correctly assembled plasmids) denotes the number of assembled plasmids matching a reference plasmid with *>* 80% sequence identity and *>* 90% reciprocal coverage. *FP* is the number of assembled plasmids that do not meet these criteria with any reference plasmid. *FN* is the number of reference plasmids lacking any corresponding assembled plasmid meeting these criteria. If multiple assembled plasmids match the same reference, only the best-matching one is counted as a *TP* .

### Performance on simulated metagenomes

We generated seven simulated metagenomes of increasing complexity, following the methodology in [18]. See Table 1 for details. For each dataset, 10 to 500 bacterial chromosomes along with their associated plasmids were selected as reference genomes. SCAPP provides a curated list of 145 bacterial strains commonly found in the human gut. For each simulation, we first selected reference bacterial genomes from this list. When additional genomes were required to reach the target reference count, we supplemented the selection with bacterial genomes randomly chosen from NCBI. Since plasmids that coexist with bacterial chromosomes are typically long, each simulation was further supplemented with a fixed number of shorter (*<* 10kb) plasmids, randomly selected from NCBI. Specifically, 5, 15, 50, 50, 100, 150, and 200 short plasmids were added in Sim1, …, Sim7, respectively. Genome abundances and plasmid copy numbers were assigned based on realistic distributions, as implemented in SCAPP (see details in the Supplementary Materials). To align with empirical observations, our simulation ensured that shorter plasmids have higher copy numbers [14]. Paired-end reads were simulated from the reference genomes at the specified abundances using InSilicoSeq [9].

**Table 1.**
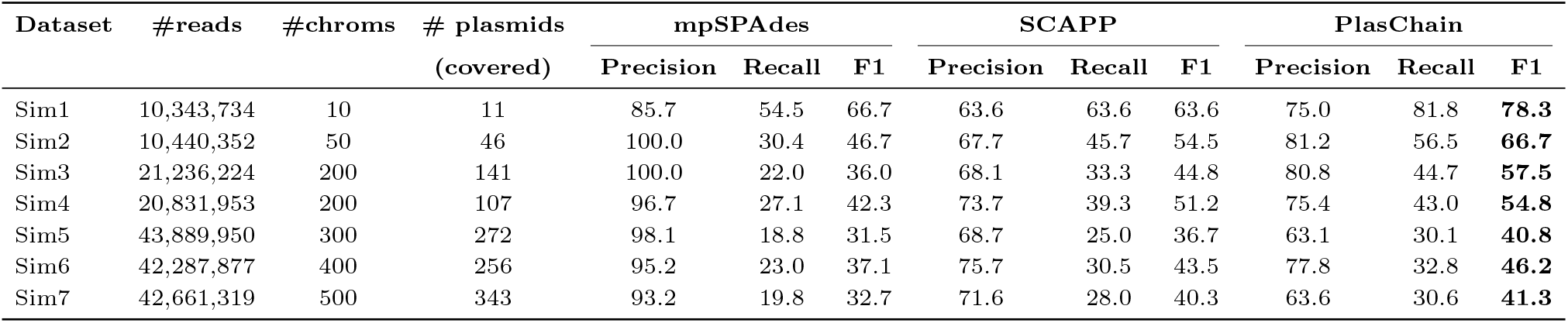
Data characteristics and assembler performance on simulated metagenomes. #chroms: number of bacterial chromosomes included in the simulated dataset. #plasmids (covered): number of plasmids included that are covered by simulated reads and can potentially be assembled. # reads: number of reads simulated. Precision, recall, and F1 scores are reported as percentages. For each simulated dataset, the highest F1 score is highlighted in bold.

| Dataset | #reads | #chroms | # plasmids<br>(covered) | mpSPAdes |  |  | SCAPP |  |  | PlasChain |  |  |
| --- | --- | --- | --- | --- | --- | --- | --- | --- | --- | --- | --- | --- |
|  |  |  |  | Precision | Recall | F1 | Precision | Recall | F1 | Precision | Recall | F1 |
| Sim1 | 10,343,734 | 10 | 11 | 85.7 | 54.5 | 66.7 | 63.6 | 63.6 | 63.6 | 75.0 | 81.8 | **78.3** |
| Sim2 | 10,440,352 | 50 | 46 | 100.0 | 30.4 | 46.7 | 67.7 | 45.7 | 54.5 | 81.2 | 56.5 | **66.7** |
| Sim3 | 21,236,224 | 200 | 141 | 100.0 | 22.0 | 36.0 | 68.1 | 33.3 | 44.8 | 80.8 | 44.7 | **57.5** |
| Sim4 | 20,831,953 | 200 | 107 | 96.7 | 27.1 | 42.3 | 73.7 | 39.3 | 51.2 | 75.4 | 43.0 | **54.8** |
| Sim5 | 43,889,950 | 300 | 272 | 98.1 | 18.8 | 31.5 | 68.7 | 25.0 | 36.7 | 63.1 | 30.1 | **40.8** |
| Sim6 | 42,287,877 | 400 | 256 | 95.2 | 23.0 | 37.1 | 75.7 | 30.5 | 43.5 | 77.8 | 32.8 | **46.2** |
| Sim7 | 42,661,319 | 500 | 343 | 93.2 | 19.8 | 32.7 | 71.6 | 28.0 | 40.3 | 63.6 | 30.6 | **41.3** |

As shown in Table 1, PlasChain outperforms SCAPP in terms of both precision and recall across nearly all tests, leading to higher F1 scores. In comparison with mpSPAdes, PlasChain achieves higher recall, while mpSPAdes demonstrates higher precision. Overall, PlasChain achieves the highest F1 scores across all test scenarios.

To further assess PlasChain’s capability in identifying long plasmids, we separately calculated precision, recall, and F1 scores for short (*<* 10kb) and long plasmids (see Table S1 in the Supplementary Materials). Figure 5 shows that all tested assemblers perform substantially better on short plasmids than on long plasmids, highlighting the greater difficulty of recovering long plasmids from short-read metagenomic assemblies. By leveraging contig path information and a cycle-merging strategy, PlasChain achieves the best overall performance on long plasmid assembly across all tested datasets. mpSPAdes outperforms SCAPP in assembling long plasmids, but it shows substantially poorer performance on short plasmid assemblies. In contrast, PlasChain provides a modest improvement over SCAPP for more than half of the short plasmid assemblies, likely due to its more stringent filtering criteria for selecting confident plasmid assemblies in the final step.

**Fig. 5.**
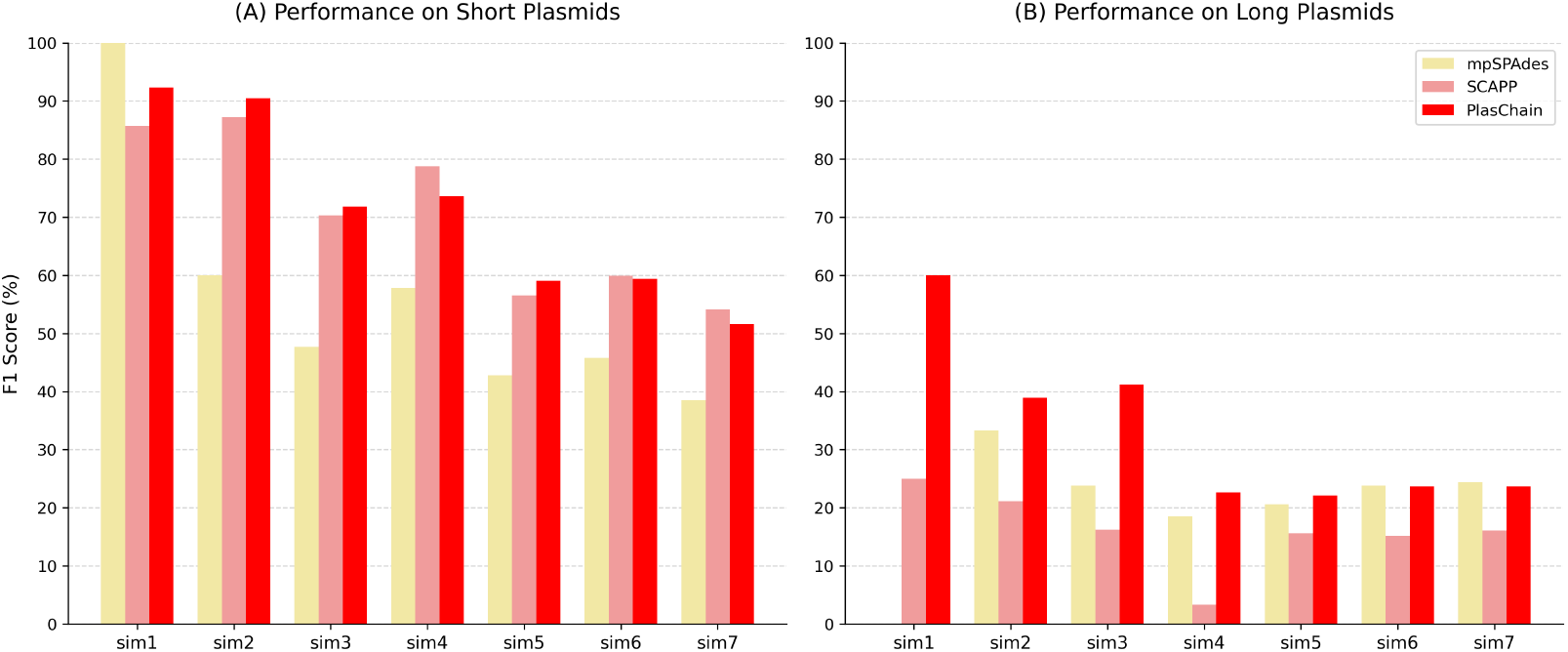
Performance in assembling long and short plasmids from simulated metagenomes.

### Performance on CAMI2 datasets

To further evaluate the performance of PlasChain in complex and realistic metagenomic environments, we benchmarked it against SCAPP and mpSPAdes on three simulated datasets from the CAMI2 challenge [8] corresponding to Airways, Skin, and Urogenital samples. These datasets are widely used for benchmarking metagenomic assembly tools [19]. They were generated from Human Microbiome Project (HMP) mock communities using the CAMISIM framework [7]. Simulated reads and the corresponding reference genomes are publicly available at https://erda.ku.dk/archives/826fe4d8889f88db2ec20058f9eaa015/published-archive.html.

Table 2 summarizes the performance of mpSPAdes, SCAPP, and PlasChain on the three CAMI2 datasets. Overall, PlasChain achieved the highest F1 score across all three environments, outperforming the other methods by 6-9 percentage points. These improvements were primarily driven by substantially higher recall while maintaining good precision.

**Table 2.**
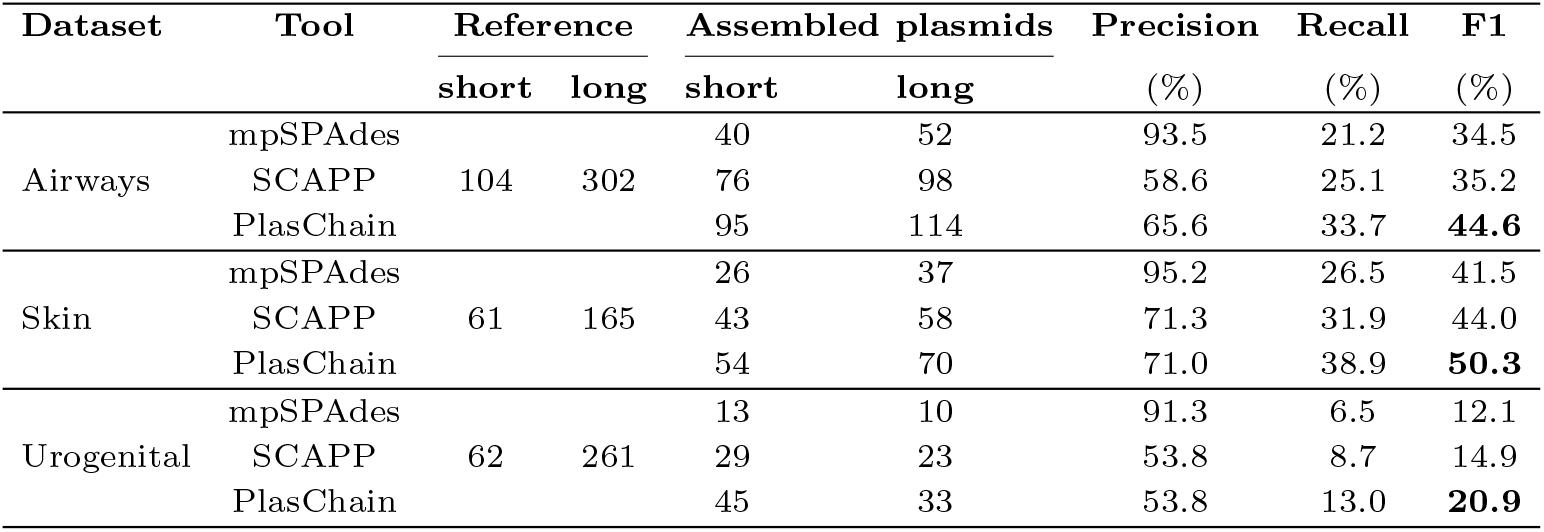
Performance on CAMI2 datasets. The columns “Reference” and “Assembled plasmids” report the number of reference plasmids identified by mapping to source genomes and the number of plasmids assembled by the tested assemblers, respectively. For each dataset, the highest F1 score is highlighted in bold.

| Dataset | Tool | Reference |  | Assembled plasmids |  | Precision | Recall | F1 |
| --- | --- | --- | --- | --- | --- | --- | --- | --- |
|  |  | short | long | short | long | (%) | (%) | (%) |
| Airways | mpSPAdes |  |  | 40 | 52 | 93.5 | 21.2 | 34.5 |
|  | SCAPP | 104 | 302 | 76 | 98 | 58.6 | 25.1 | 35.2 |
|  | PlasChain |  |  | 95 | 114 | 65.6 | 33.7 | **44.6** |
| Skin | mpSPAdes |  |  | 26 | 37 | 95.2 | 26.5 | 41.5 |
|  | SCAPP | 61 | 165 | 43 | 58 | 71.3 | 31.9 | 44.0 |
|  | PlasChain |  |  | 54 | 70 | 71.0 | 38.9 | **50.3** |
| Urogenital | mpSPAdes |  |  | 13 | 10 | 91.3 | 6.5 | 12.1 |
|  | SCAPP | 62 | 261 | 29 | 23 | 53.8 | 8.7 | 14.9 |
|  | PlasChain |  |  | 45 | 33 | 53.8 | 13.0 | **20.9** |

mpSPAdes consistently achieved the highest precision (91.3– 95.2%) across all datasets, reflecting its conservative assembly strategy. However, this conservative approach resulted in markedly lower recall (6.5–26.5%), indicating that many reference plasmids were not recovered. In contrast, PlasChain achieved a substantially better balance between precision and recall. Compared with SCAPP, PlasChain consistently increased recall by 4.3–8.6 percentage points while maintaining similar precision, leading to the highest overall F1 scores. Furthermore, Supplementary Figure S2 and Table S2 show that these improvements were observed for both short and long plasmids, demonstrating that the proposed contig-path integration and cycle-merging strategy benefits plasmid assembly across a broad range of plasmid lengths.

### Performance on real-world metagenomic datasets

To evaluate the assemblers on real-world data, we collected six metagenomic and plasmidomic datasets: (1-3) INFANT 32, INFANT 302, INFANT 332: Infant fecal samples of taken at day 32, 302, and 332 after birth. Data was taken from [25], NCBI accession IDs: SRR28256440, SRR28256441, and SRR28256439. (4) Human Gut Plasmidome (HGP): plasmidome of a healthy adult’s gut microbiome [18], NCBI accession ID: SRR11038083. Plasmidome sequencing enriches plasmid-derived DNA, thereby increasing the chance of recovering plasmids while reducing interference from chromosomal sequences. (5) MARINE: A metagenome from marine sediments collected near active hydrothermal vents in the Atlantic Ocean [24], NCBI accession ID: SRX684858. (6) LAKE: A metagenome derived from a lake in India contaminated with fluoroquinolone antibiotics due to industrial discharge [2], NCBI accession ID: ERS433966.

Since the true plasmid content of real sequencing datasets is unknown, we identify potentially present plasmids by searching for matches in the PLSDB database [15] and use these as the gold standard for evaluation. Because all tested assemblers rely on metaSPAdes as the underlying assembler, plasmid reconstruction is performed from its contig output. Therefore, all contigs generated by metaSPAdes are aligned to plasmids in PLSDB using BLAST. A contig is defined as *plasmidic* if it aligns to a plasmid genome with *>* 85% sequence identity and the alignment covers *>* 85% of the contig length. Plasmids in PLSDB for which *>* 90% of their genome is covered by plasmidic contigs are selected as the gold-standard reference set. If multiple PLSDB plasmids are supported by the same set of plasmidic contigs, only the plasmid with the highest genome coverage is retained. The number of identified plasmids for each dataset is reported in Table 3.

**Table 3.** Performance on real metagenomic datasets. The columns “PLSDB plasmids” and “Assembled plasmids” report the number of reference plasmids identified by mapping to PLSDB and the number of plasmids assembled by each tested method, respectively. For each dataset, the highest F1 score is highlighted in bold.

| Dataset | Tool | PLSDB plasmids |  | Assembled plasmids |  | Precision | Recall | F1 |
| --- | --- | --- | --- | --- | --- | --- | --- | --- |
|  |  | short | long | short | long | (%) | (%) | (%) |
| INFANT_32 | mpSPAdes |  |  | 3 | 5 | 12.5 | 10.0 | 11.1 |
|  | SCAPP | 9 | 1 | 4 | 3 | 57.1 | 40.0 | 47.1 |
|  | PlasChain |  |  | 4 | 4 | 62.5 | 50.0 | **55.6** |
| INFANT_302 | mpSPAdes |  |  | 8 | 2 | 50.0 | 21.7 | 30.3 |
|  | SCAPP | 20 | 3 | 11 | 2 | 46.2 | 26.1 | 33.3 |
|  | PlasChain |  |  | 12 | 2 | 57.1 | 34.8 | **43.2** |
| INFANT_332 | mpSPAdes |  |  | 12 | 3 | 46.7 | 21.9 | 29.8 |
|  | SCAPP | 23 | 9 | 17 | 5 | 45.5 | 31.2 | 37.0 |
|  | PlasChain |  |  | 17 | 6 | 47.8 | 34.4 | **40.0** |
| HGP | mpSPAdes |  |  | 48 | 0 | 25.0 | 27.3 | 26.1 |
|  | SCAPP | 43 | 1 | 84 | 0 | 20.2 | 38.6 | 26.6 |
|  | PlasChain |  |  | 85 | 0 | 21.2 | 40.9 | **27.9** |
| MARINE | mpSPAdes |  |  | 9 | 33 | 2.4 | 25.0 | **4.3** |
|  | SCAPP | 4 | 0 | 46 | 56 | 1.0 | 25.0 | 1.9 |
|  | PlasChain |  |  | 47 | 61 | 0.9 | 25.0 | 1.8 |
| LAKE | mpSPAdes |  |  | 395 | 14 | 2.4 | 1.6 | 1.9 |
|  | SCAPP | 626 | 5 | 1,003 | 14 | 2.2 | 3.5 | 2.7 |
|  | PlasChain |  |  | 1,096 | 20 | 2.2 | 4.0 | **2.9** |

Due to their complexity, the performance of all tested assemblers is lower on some of the real sequencing datasets (Table 3). Still, PlasChain consistently outperforms SCAPP in both precision and recall across nearly all datasets. Although all assemblers exhibit low recall, PlasChain achieves the highest recall in all datasets. In terms of precision, PlasChain yields the best performance in three datasets, while mpSPAdes performs best in the other three. Overall, PlasChain has the best F1 score in five of the six datasets, with the exception of MARINE. This may be explained by the absence of long gold standard plasmids in MARINE, since PlasChain is designed for better detection of long plasmids. Supplementary Table S3 further reveals that, although all assemblers reconstructed some long plasmids from five of the six real datasets, PlasChain was the only method that reconstructed plasmids matching reference genomes in the PLSDB database. In contrast, none of the long plasmids reconstructed by the other assemblers matched any reference genome in PLSDB.

In the MARINE dataset, only four reference plasmids were identified using PLSDB, all of which are short. However, all tested assemblers reconstructed a much larger number of plasmids from this dataset, and more than half are long plasmids. This discrepancy raises the possibility that the MARINE environment may harbor numerous long novel plasmids that are not yet included in PLSDB. To further evaluate this hypothesis, we used geNomad [5], a state-of-the-art tool for identifying plasmids and viruses from metagenome assemblies that is well suited for detecting novel species. The results of applying geNomad to each assembly are summarized in Table 4, showing that PlasChain substantially outperforms both mpSPAdes and SCAPP in terms of the number of plasmids identified by geNomad, their length, and gene annotations. Among the plasmids assembled by PlasChain, ten were verified by geNomad, including one exceeding 310 kb in length, potentially representing a megaplasmid. Notably, six of these ten putative plasmids contain conjugation-associated genes or antimicrobial resistance (AMR) genes. In contrast, mpSPAdes and SCAPP recover fewer and shorter plasmids, often lacking key functional markers.

**Table 4.**
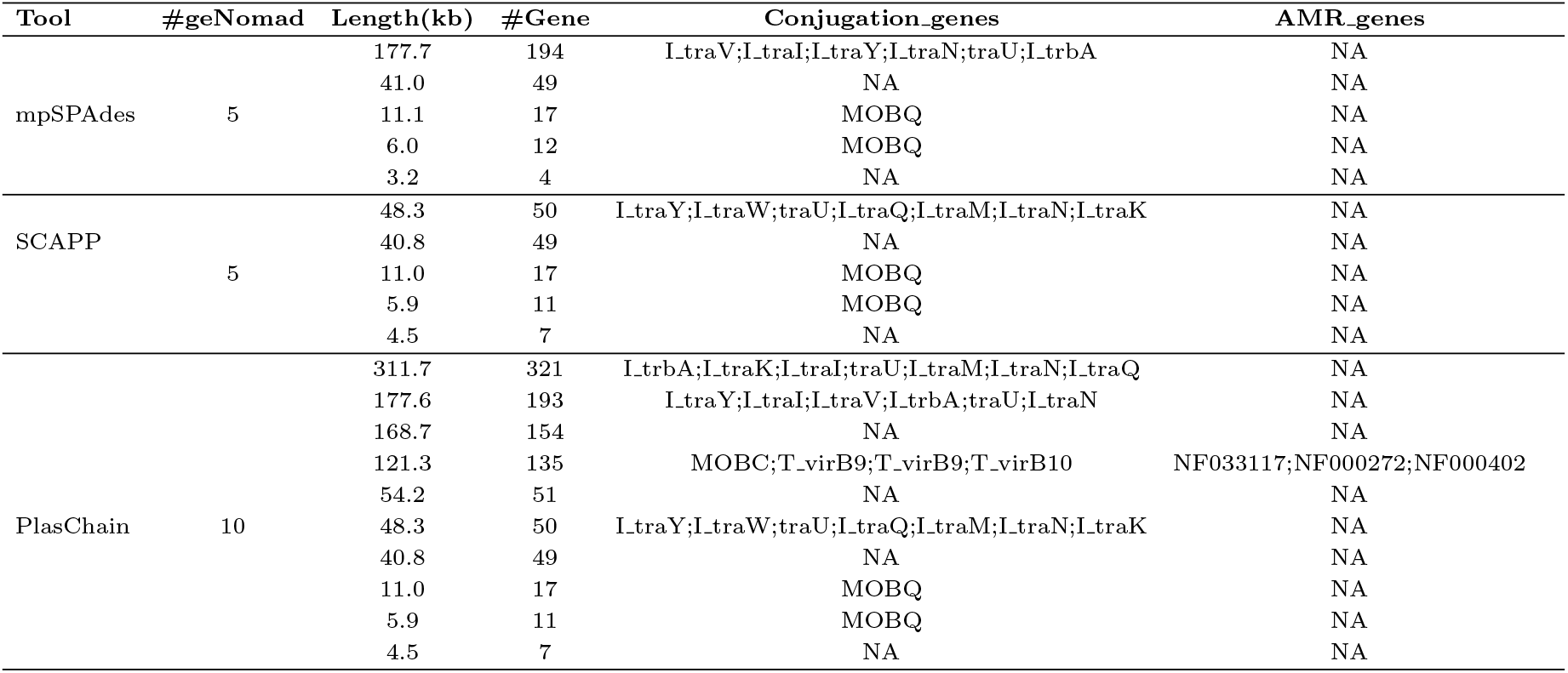
Identification of novel plasmids from the MARINE dataset. #geNomad: number of plasmids assembled by each method that were identified as plasmids by geNomad. For each such plasmid, its length, gene count, presence of conjugation-associated genes, and antimicrobial resistance (AMR) genes are listed as supporting evidence for plasmid identification.

### Resource usage and availability

The computational resources used by each assembler and each dataset is summarized in Table 5. All experiments were performed on a server equipped with 32 CPU cores (3.1 GHz) and 503 GB RAM. Both SCAPP and PlasChain operate on the assembly graph generated by metaSPAdes. The graph construction represents a standard preprocessing step in metagenomic assembly rather than an additional computational cost specific to either method.

**Table 5.** Runtime and peak memory usage of plasmid assemblers. Total execution time (minutes) and peak memory consumption (GB) are reported for mpSPAdes, SCAPP, PlasChain, and metaSPAdes on simulated and real-world metagenomic datasets.

| Sample | Time (minutes) |  |  |  | Memory (GB) |  |  |  |
| --- | --- | --- | --- | --- | --- | --- | --- | --- |
|  | mpSPAdes | SCAPP | PlasChain | metaSPAdes | mpSPAdes | SCAPP | PlasChain | metaSPAdes |
| Sim1 | 16 | 19 | 23 | 13 | 15.7 | 1.7 | 4.2 | 15.6 |
| Sim2 | 26 | 38 | 48 | 22 | 16.1 | 12.8 | 12.6 | 16.2 |
| Sim3 | 68 | 112 | 136 | 60 | 39.7 | 12.8 | 12.8 | 39.4 |
| Sim4 | 76 | 128 | 138 | 73 | 41.3 | 12.8 | 12.8 | 41.4 |
| Sim5 | 152 | 310 | 805 | 145 | 75.0 | 12.8 | 12.8 | 75.0 |
| Sim6 | 209 | 366 | 530 | 226 | 73.0 | 12.9 | 12.9 | 72.9 |
| Sim7 | 451 | 413 | 516 | 246 | 83.6 | 12.9 | 12.9 | 83.8 |
| HGP | 183 | 213 | 214 | 178 | 18.3 | 18.6 | 19.0 | 18.3 |
| MARINE | 288 | 348 | 358 | 267 | 71.6 | 12.9 | 12.9 | 72.0 |
| LAKE | 113 | 161 | 86 | 127 | 39.2 | 20.0 | 20.0 | 39.1 |
| INFANT32 | 255 | 304 | 423 | 266 | 35.6 | 12.5 | 12.5 | 37.7 |
| INFANT302 | 217 | 268 | 373 | 236 | 20.3 | 2.3 | 7.5 | 21.5 |
| INFANT332 | 173 | 226 | 270 | 197 | 23.7 | 12.7 | 12.6 | 23.7 |

In terms of runtime, PlasChain generally required more time than SCAPP because of its additional contig-path processing and exhaustive cycle-merging procedure. Runtime ranged from 23 minutes on Sim1 to 805 minutes on Sim5, depending on dataset complexity. Nevertheless, time difference varied considerably across datasets. On the LAKE dataset, for example, PlasChain required only 86 minutes, outperforming both SCAPP (161 minutes) and mpSPAdes (113 minutes), suggesting that the proposed graph simplification strategy can substantially accelerate processing on certain highly tangled assembly graphs.

Peak memory usage was dominated by the assembly graph construction performed by metaSPAdes. Consequently, PlasChain and SCAPP exhibited nearly identical peak memory footprints across all datasets, since both of them operate on the assembly graph generated by metaSPAdes. Likewise, mpSPAdes showed a similar peak memory footprint because it is also built on top of metaSPAdes. These results indicate that the improved plasmid recovery achieved by PlasChain comes with only a modest runtime overhead and with virtually no additional memory consumption.

PlasChain is open source and is freely available at https://github.com/SDU-ACG-Lab/PlasChain.

## Discussion and conclusions

Although the critical role of plasmids in the dissemination of antibiotic resistance is widely recognized, methods for recovering complete plasmid sequences from metagenomic samples remain limited. In this work, we introduced PlasChain, a novel plasmid assembler designed to improve plasmid recovery from metagenomic assemblies. By integrating contig path information and a cycle-merging strategy, PlasChain exhibits a pronounced advantage in assembling long plasmids on both simulated and real datasets, while maintaining competitive performance on short plasmids. We note that while the cycle-merging step may occasionally merge distinct plasmids that share repetitive sequences, Supplementary Figure S1 indicates that this strategy improves overall performance across most evaluated datasets.

All evaluated assemblers, PlasChain included, show reduced accuracy on real metagenomic datasets due to their inherent complexity. However, it is important to note that the current gold-standard evaluation, which relies on the PLSDB database, likely underestimates the true plasmid content of these samples. Our results on a metagenome sample from a less explored environment (MARINE) suggest that PlasChain can reconstruct previously uncharacterized plasmids, including megaplasmids that are typically underrepresented in existing databases. A robust and broadly accepted framework for defining gold standards in real-world metagenomic analyses is still lacking and remains an important area for future work.

In addition, although the majority of plasmids are circular, linear plasmids have been reported in several bacterial species [11]. However, existing plasmid assemblers, including plasChain, are primarily designed for circular plasmids, leaving the identification and reconstruction of linear plasmids an important but still unexplored problem.

Despite incorporating additional contig path information, the performance gains of PlasChain over SCAPP and mpSPAdes remain moderate, largely due to the inherent difficulty of reconstructing plasmids from short-read sequencing data. Advances in sequencing technologies may help address these limitations. In particular, third-generation long-read sequencing platforms offer substantially longer reads that can facilitate the assembly of large plasmids, although their higher error rates introduce new computational challenges. Our future work will focus on developing methods tailored to long-read data, as well as hybrid strategies that integrate both long and short reads to improve plasmid recovery from metagenomic samples.

## Declarations

## Acknowledgements

We thank members of the SDU-ACG-Lab for their help and advice: Xiaolin Li, Xun Ding, and Lin Gao.

## Author’s contributions

SF and HS contributed equally to this work. SF and HS developed and implemented the PlasChain algorithm, conducted benchmark experiments, analyzed the results, and drafted the manuscript. LR conceived and designed the study. Both RS and LP supervised the development and evaluation of the method, provided critical guidance throughout the project, and revised the manuscript. All authors read and approved the final version of the manuscript.

## Availability of data and materials

The source code and testing data for PlasChain are publicly available at https://github.com/SDU-ACG-Lab/PlasChain. The sequencing reads of the CAMI2 datasets are publicly available through the CAMI2 data repository. The sequencing reads of the INFANT 32, INFANT 302, and INFANT 332 datasets are available from NCBI under the accession IDs SRR28256440, SRR28256441, and SRR28256439, respectively. The sequencing reads of the Human Gut Plasmidome (HGP), MARINE, and LAKE datasets are available from NCBI under the accession IDs SRR11038083, SRX684858, and ERS433966, respectively.

## Ethics approval and consent to participate

Not applicable.

## Consent for publication

Not applicable.

