## Supplementary Information for "PlasChain: an algorithm for improving long plasmid reconstruction from metagenome assemblies"

#### 1 Definition of discounted coverage

To evaluate coverage consistency within a candidate cycle identified during the cycle-peeling step, we define the *discounted coverage* for each node in the cycle as follows. Let $u$ be a node belonging to a candidate cycle $C$, and let $\mathcal{N}(u)$ denote the set of neighbors of $u$ in the assembly graph. We further define $\mathcal{N}_{\mathrm{C}}(u) \subseteq \mathcal{N}(u)$ as the subset of neighbors that also belong to cycle $C$. Let $cov(u)$ denote the sequencing coverage of node $u$ as reported by the assembler. The discounted coverage of node $u$, denoted $cov'(u)$, is defined as:

$$cov'(u) = cov(u) \times \frac{\sum_{v \in \mathcal{N}_C(u)} \mathrm{cov}(v)}{\sum_{v \in \mathcal{N}(u)} \mathrm{cov}(v)}. \tag{1}$$

This formulation adjusts the raw coverage of node $u$ by weighting it according to the proportion of neighboring coverage contributed by nodes within the candidate cycle $C$. As a result, branch nodes that are shared by multiple candidate cycles or connected to alternative graph paths receive a reduced effective coverage, reflecting the fact that part of their sequencing coverage may originate from sequences outside the current cycle. By discounting these ambiguous nodes, the resulting discounted coverage more accurately captures the intrinsic coverage variation of the candidate cycle, leading to a more reliable assessment of plasmid candidates.

#### 2 Generation of simulated metagenomes

Seven simulated metagenomic datasets of increasing complexity were generated following the simulation strategy described in SCAPP. For each dataset, 10 to 500 bacterial chromosomes and their associated plasmids were selected as reference genomes. Because plasmids coexisting with bacterial chromosomes are generally long, each simulated community was supplemented with a fixed number of short plasmids (< 10kb), randomly selected from NCBI RefSeq database. Specifically, 5, 15, 50, 50, 100, 150, and 200 short plasmids were added to Sim1, ..., Sim7, respectively.

Paired-end Illumina reads were then simulated separately for bacterial chromosomes and plasmids using InSilicoSeq. The relative abundance of bacterial chromosomes were sampled from a log-normal distribution. Plasmid copy numbers were independently sampled from a geometric distribution with parameter $p$, where shorter plasmids were assigned higher copy numbers to reflect their higher abundance in natural microbial communities. The parameter $p$ was defined as a function of plasmid length $L$:

$$p = \begin{cases} \log_{10}(L)/30, & 1\text{kb} \leq L < 10\text{kb} \\ \log_{10}(L)/20, & 10\text{kb} \leq L < 100\text{kb} \\ \log_{10}(L)/10, & 100\text{kb} \leq L < 1\text{Mb} \\ 1, & L \geq 1\text{Mb} \end{cases} \tag{2}$$

Here, $p$ denotes the parameter of the geometric distribution for a plasmid of length $L$. A total of 5 million paired-end chromosome reads were generated for Sim1 and Sim2, 10 million for Sim3 and Sim4, and 20 million for Sim5, Sim6, and Sim7. The number of reads generated for each plasmid was determined by its copy number and the sequencing depth of its host chromosome. Reads from plasmids and chromosomes were simulated using the same sequencing error model and subsequently combined to generate the final metagenomic datasets.

### 3 Parameter sensitivity analysis of the cycle-merging strategy

To evaluate the robustness of the cycle-merging strategy, we generated five independent validation metagenomic datasets of increasing complexity following the simulation procedure described in Section 2. Specifically, the datasets contained 50–400 bacterial chromosomes and 15–200 plasmids, respectively. We then assessed the performance of PlasChain under different combinations of the coverage similarity and tetranucleotide composition similarity thresholds used during cycle merging (Figure S1).

Compared with the baseline without cycle merging, incorporating the cycle-merging step consistently improved plasmid assembly performance across all five validation datasets and for the vast majority of parameter combinations. Although the cycle-merging strategy may occasionally merge distinct plasmids that share repetitive sequences, the overall improvement in F1 score demonstrates that its benefits substantially outweigh these rare incorrect merges.

Overall, the cycle-merging strategy is robust to the choice of both parameters. Varying the coverage similarity threshold from 0.05 to 0.30 changed the F1 score by less than 2.5 percentage points across all validation datasets. Performance generally improved as the threshold increased and remained stable for thresholds between 0.20 and 0.30, with a threshold of 0.20 providing consistently strong performance across datasets of different complexity. Likewise, varying the tetranucleotide composition similarity threshold from 0.70 to 0.95 resulted in only minor performance fluctuations. The only noticeable exception occurred at the most stringent threshold (0.95), which slightly reduced the F1 score on the simplest validation dataset (Figure S1 (a)). Based on these results, PlasChain adopts default thresholds of 0.20 for coverage similarity and 0.90 for tetranucleotide composition similarity, which provide a favorable balance between assembly accuracy and robustness across diverse metagenomic communities.

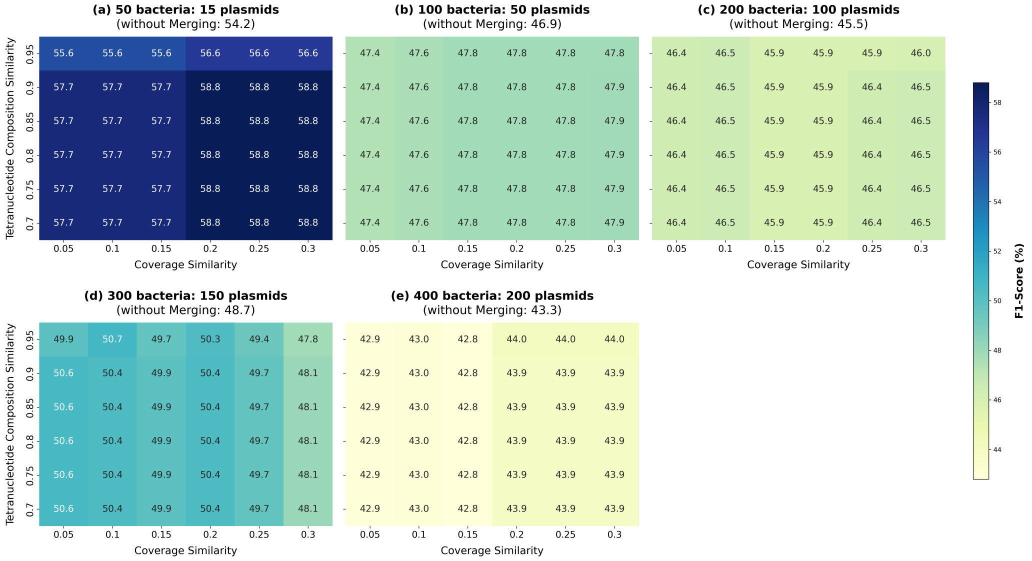


Figure S1: Parameter sensitivity analysis of the cycle-merging strategy across simulated metagenomic datasets of increasing complexity. Heatmaps show the F1 score (%) obtained under different combinations of the tetranucleotide composition similarity and coverage similarity thresholds. Panels (a-e) correspond to datasets containing 50, 100, 200, 300, and 400 bacterial chromosomes, respectively. The baseline F1 score without cycle merging is shown in parentheses above each heatmap. Warmer colors indicate higher F1 scores.

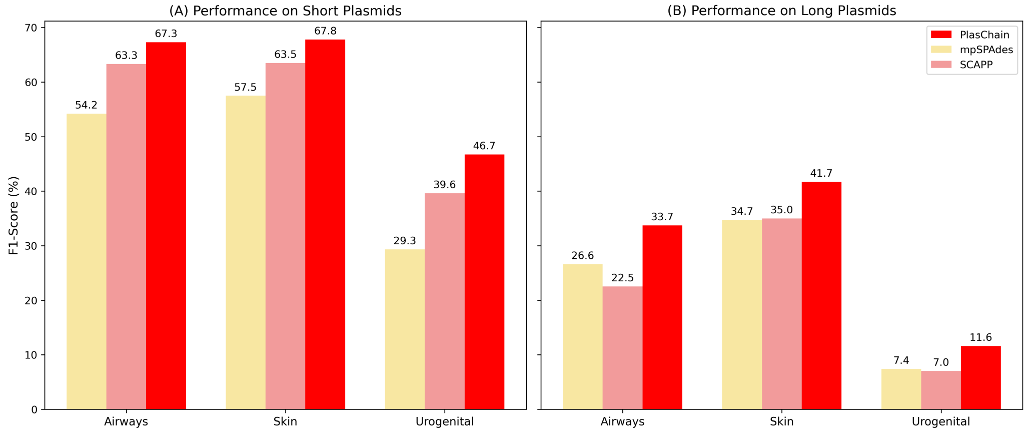


Figure S2: Assembly performance on short and long plasmids in the three CAMI2 simulated metagenomic datasets. F1 scores (%) achieved by PlasChain, SCAPP, and mpSPAdes are shown separately for (A) short plasmids and (B) long plasmids in the Airways, Skin, and Urogenital datasets.

Table S1: Performance of mpSPAdes, SCAPP, and PlasChain on simulated metagenomes. #plasmids (covered): number of plasmids included that are covered by simulated reads and can potentially be assembled. Precision, recall, and F1 score are reported separately for short and long plasmids as percentages. Red, blue, and orange numbers indicate the highest overall F1 score, the highest F1 score for short plasmids, and the highest F1 score for long plasmids, respectively.

| Dataset | # plasmids | mpSPAdes | | | SCAPP | | | PlasChain | | |
|---|---|---|---|---|---|---|---|---|---|---|
| | (covered) | Precision | Recall | **F1 score** | Precision | Recall | **F1 score** | Precision | Recall | **F1 score** |
| sim1 | 11 | 85.7 | 54.5 | 66.7 | 63.6 | 63.6 | 63.6 | 75.0 | 81.8 | **78.3** |
| sim1 (short) | 6 | 100.0 | 100.0 | **100.0** | 75.0 | 100.0 | 85.7 | 85.7 | 100.0 | 92.3 |
| sim1 (long) | 5 | 0.0 | 0.0 | 0.0 | 33.3 | 20.0 | 25.0 | 60.0 | 60.0 | **60.0** |
| sim2 | 46 | 100.0 | 30.4 | 46.7 | 67.7 | 45.7 | 54.5 | 81.2 | 56.5 | **66.7** |
| sim2 (short) | 21 | 100.0 | 42.9 | 60.0 | 94.4 | 81.0 | 87.2 | 90.0 | 90.5 | **90.5** |
| sim2 (long) | 25 | 100.0 | 20.0 | 33.3 | 30.8 | 16.0 | 21.1 | 63.6 | 28.0 | **38.9** |
| sim3 | 141 | 100.0 | 22.0 | 36.0 | 68.1 | 33.0 | 44.8 | 80.8 | 44.7 | **57.5** |
| sim3 (short) | 67 | 100.0 | 31.3 | 47.7 | 88.6 | 58.2 | 70.3 | 84.0 | 62.7 | **71.8** |
| sim3 (long) | 74 | 100.0 | 13.5 | 23.8 | 32.0 | 10.8 | 16.2 | 75.0 | 28.4 | **41.2** |
| sim4 | 107 | 96.7 | 27.1 | 42.3 | 73.7 | 39.3 | 51.2 | 75.4 | 43.0 | **54.8** |
| sim4 (short) | 59 | 100.0 | 40.7 | 57.8 | 91.1 | 69.5 | **78.8** | 83.0 | 66.1 | 73.6 |
| sim4 (long) | 48 | 83.3 | 10.4 | 18.5 | 8.3 | 2.1 | 3.3 | 50.0 | 14.6 | **22.6** |
| sim5 | 272 | 98.1 | 18.8 | 31.5 | 68.7 | 25.0 | 36.7 | 63.1 | 30.1 | **40.8** |
| sim5 (short) | 124 | 97.1 | 27.4 | 42.8 | 80.6 | 43.5 | 56.5 | 75.9 | 48.4 | **59.1** |
| sim5 (long) | 148 | 100.0 | 11.5 | 20.6 | 43.8 | 9.5 | 15.6 | 43.1 | 14.9 | **22.1** |
| sim6 | 256 | 95.2 | 23.0 | 37.1 | 75.7 | 30.5 | 43.5 | 77.8 | 32.8 | **46.2** |
| sim6 (short) | 147 | 97.8 | 29.9 | 45.8 | 85.0 | 46.3 | **59.9** | 82.9 | 46.3 | 59.4 |
| sim6 (long) | 109 | 88.2 | 13.8 | **23.8** | 43.5 | 9.2 | 15.2 | 61.5 | 14.7 | 23.7 |
| sim7 | 343 | 93.2 | 19.8 | 32.7 | 71.6 | 28.0 | 40.3 | 63.6 | 30.6 | **41.3** |
| sim7 (short) | 197 | 100.0 | 23.9 | 38.5 | 77.4 | 41.6 | **54.1** | 66.4 | 42.1 | 51.6 |
| sim7 (long) | 146 | 80.8 | 14.4 | **24.4** | 50.0 | 9.6 | 16.1 | 55.0 | 15.1 | 23.7 |

Table S2: Performance of mpSPAdes, SCAPP, and PlasChain on CAMI2 dataset.The 'Covered' column indicates the total number of reference plasmids in the dataset, whereas the 'Assembly' column represents the number of plasmids reconstructed by each tool. Precision, recall, and F1 score are reported as percentages and categorized by plasmid length (all, short, and long). Red indicates the highest overall F1 score, blue highlights the optimal F1 score for short plasmids, and orange denotes the best F1 score for long plasmids.

| Simulation | Tool | Category | Covered | Assembly | Precision | Recall | F1 Score |
|---|---|---|---|---|---|---|---|
| **Airways** | **PlasChain** | All | 406 | 209 | 65.6 | 33.7 | **44.6** |
| | | Short | 104 | 95 | 70.5 | 64.4 | **67.3** |
| | | Long | 302 | 114 | 61.4 | 23.2 | **33.7** |
| | **mpSPAdes** | All | 406 | 92 | 93.5 | 21.2 | 34.5 |
| | | Short | 104 | 40 | 97.5 | 37.5 | 54.2 |
| | | Long | 302 | 52 | 90.4 | 15.6 | 26.6 |
| | **SCAPP** | All | 406 | 174 | 58.6 | 25.1 | 35.2 |
| | | Short | 104 | 76 | 75.0 | 54.8 | 63.3 |
| | | Long | 302 | 98 | 45.9 | 14.9 | 22.5 |
| **Skin** | **PlasChain** | All | 226 | 124 | 71.0 | 38.9 | **50.3** |
| | | Short | 61 | 54 | 72.2 | 63.9 | **67.8** |
| | | Long | 165 | 70 | 70.0 | 29.7 | **41.7** |
| | **mpSPAdes** | All | 226 | 63 | 95.2 | 26.5 | 41.5 |
| | | Short | 61 | 26 | 96.2 | 41.0 | 57.5 |
| | | Long | 165 | 37 | 94.6 | 21.2 | 34.7 |
| | **SCAPP** | All | 226 | 101 | 71.3 | 31.9 | 44.0 |
| | | Short | 61 | 43 | 76.7 | 54.1 | 63.5 |
| | | Long | 165 | 58 | 67.2 | 23.6 | 35.0 |
| **Urogenital** | **PlasChain** | All | 323 | 78 | 53.8 | 13.0 | **20.9** |
| | | Short | 62 | 45 | 55.6 | 40.3 | **46.7** |
| | | Long | 261 | 33 | 51.5 | 6.5 | **11.6** |
| | **mpSPAdes** | All | 323 | 23 | 91.3 | 6.5 | 12.1 |
| | | Short | 62 | 13 | 84.6 | 17.7 | 29.3 |
| | | Long | 261 | 10 | 100 | 3.8 | 7.4 |
| | **SCAPP** | All | 323 | 52 | 53.8 | 8.7 | 14.9 |
| | | Short | 62 | 29 | 62.1 | 29.0 | 39.6 |
| | | Long | 261 | 23 | 43.5 | 3.8 | 7.0 |

Table S3: Performance of tested assemblers on six real datasets. "PLSDB plasmids" reports the number of reference plasmids identified through mapping to PLSDB. Precision, recall, and F1 score are reported separately for short and long plasmids as percentages. Red indicates the highest overall F1 score, blue highlights the best F1 score for short plasmids, and orange highlights the best F1 score for long plasmids.

| Dataset | PLSDB plasmids | mpSPAdes | | | SCAPP | | | PlasChain | | |
|---|---|---|---|---|---|---|---|---|---|---|
| | | Precision | Recall | **F1 score** | Precision | Recall | **F1 score** | Precision | Recall | **F1 score** |
| INFANT32 | 10 | 12.5 | 10.0 | 11.1 | 57.1 | 40.0 | 47.1 | 62.5 | 50.0 | **55.6** |
| INFANT32 (short) | 9 | 33.3 | 11.1 | 16.7 | 100.0 | 44.4 | **61.5** | 100.0 | 44.4 | **61.5** |
| INFANT32 (long) | 1 | 0.0 | 0.0 | 0.0 | 0.0 | 0.0 | 0.0 | 25.0 | 100.0 | 40.0 |
| INFANT302 | 23 | 50.0 | 21.7 | 30.3 | 46.2 | 26.1 | 33.3 | 57.1 | 34.8 | **43.2** |
| INFANT302 (short) | 20 | 62.5 | 25.0 | 35.7 | 54.5 | 30.0 | 38.7 | 58.3 | 35.0 | **43.8** |
| INFANT302 (long) | 3 | 0.0 | 0.0 | 0.0 | 0.0 | 0.0 | 0.0 | 50.0 | 33.3 | 40.0 |
| INFANT332 | 32 | 46.7 | 21.9 | 29.8 | 45.5 | 31.2 | 37.0 | 47.8 | 34.4 | **40.0** |
| INFANT332 (short) | 23 | 58.3 | 30.4 | 40.0 | 58.8 | 43.5 | **50.0** | 58.8 | 43.5 | **50.0** |
| INFANT332 (long) | 9 | 0.0 | 0.0 | 0.0 | 0.0 | 0.0 | 0.0 | 16.7 | 11.1 | 13.3 |
| HGP | 44 | 25.0 | 27.3 | 26.1 | 20.2 | 38.6 | 26.6 | 21.2 | 40.9 | **27.9** |
| HGP (short) | 43 | 25.0 | 27.9 | 26.4 | 20.2 | 39.5 | 26.8 | 21.2 | 41.9 | **28.1** |
| HGP (long) | 1 | 0.0 | 0.0 | 0.0 | 0.0 | 0.0 | 0.0 | 0.0 | 0.0 | 0.0 |
| MARINE | 4 | 2.4 | 25.0 | **4.3** | 1.0 | 25.0 | 1.9 | 0.9 | 25.0 | 1.8 |
| MARINE (short) | 4 | 11.1 | 25.0 | **15.4** | 2.2 | 25.0 | 4.0 | 2.1 | 25.0 | 3.9 |
| MARINE (long) | 0 | 0.0 | 0.0 | 0.0 | 0.0 | 0.0 | 0.0 | 0.0 | 0.0 | 0.0 |
| LAKE | 631 | 2.4 | 1.6 | 1.9 | 2.2 | 3.5 | 2.7 | 2.2 | 4.0 | **2.9** |
| LAKE (short) | 626 | 2.5 | 1.6 | 2.0 | 2.2 | 3.5 | 2.7 | 2.3 | 4.0 | **2.9** |
| LAKE (long) | 5 | 0.0 | 0.0 | 0.0 | 0.0 | 0.0 | 0.0 | 0.0 | 0.0 | 0.0 |
